# Structural basis for ssDNA-activated NADase activity of the prokaryotic SPARTA immune system

**DOI:** 10.1101/2023.07.14.549122

**Authors:** Jun-Tao Zhang, Xin-Yang Wei, Ning Cui, Ruilin Tian, Ning Jia

## Abstract

Argonaute proteins (Agos), which use small RNAs or DNAs as guides to recognize complementary nucleic acid targets, mediate RNA silencing in eukaryotes. In prokaryotes, Agos are involved in immunity: the short prokaryotic Ago/TIR-APAZ (SPARTA) immune system triggers cell death by degrading NAD^+^ in response to invading plasmids, but its molecular mechanisms remain unknown. Here, we used cryogenic electron microscopy to determine the structures of inactive monomeric and active tetrameric *Crenotalea thermophila* SPARTA complexes, revealing mechanisms underlying SPARTA assembly, RNA-guided recognition of target single-stranded DNA (ssDNA) and subsequent SPARTA tetramerization, as well as tetramerization-dependent NADase activation. The small RNA guides Ago to recognize its ssDNA target, inducing SPARTA tetramerization via both Ago- and TIR-mediated interactions and resulting in a two-stranded, parallel, head-to-tail TIR rearrangement primed for NAD^+^ hydrolysis. Our findings thus identify the molecular basis for target ssDNA-mediated SPARTA activation, which will facilitate the development of SPARTA-based biotechnological tools.

## Main

Detection of foreign nucleic acids is a common aspect of cellular immunity across all domains of life. Argonaute proteins (Agos) use small oligonucleotides as guides for sequence-specific recognition of complementary nucleic acid targets^1, 2^. Eukaryotic Ago proteins (eAgos) play crucial roles in gene silencing pathways through RNA-guided RNA targeting^3, 4^, whereas prokaryotic Ago proteins (pAgos) function in prokaryotic immune systems against invading plasmids and viruses^1, 5–7^. eAgos have a conserved bilobed architecture, with the N-terminal lobe consisting of the N and Piwi-Argonaute-Zwille (PAZ) domains and the C-terminal lobe consisting of the Middle (MID) and P-element-induced wimpy testis (PIWI) domains^4^. The PAZ and MID domains anchor guide RNA (gRNA) at the 3′ and 5′ ends, respectively, while the PIWI domain contains an RNase H-like fold with a conserved catalytic tetrad that is responsible for cleavage of the target RNA^8^. The N domain serves as a wedge to facilitate the separation of gRNA and target RNA strands^4, 9^. The RNA-guided RNA interference ability of eAgos has made them promising tools for precise gene silencing in both basic research and therapeutic applications^10^.

In contrast to eAgos, pAgos utilize small RNA or single-stranded DNA (ssDNA) guides to recognize RNA or DNA targets^1, 5–7^. In addition, pAgos have a much more diverse domain composition, and are phylogenetically classified as long or short pAgos^1, 6, 7^. Long pAgos share a domain architecture similar to that of eAgos, whereas short pAgos contain only the MID domain and a catalytically inactive PIWI domain. Short pAgos are typically associated with analog of PAZ (APAZ)-containing proteins, which are usually fused with putative nuclease families such as Mrr, Toll/interleukin-1 receptor (TIR), and Silent information regulator 2 (SIR2)^1, 5–7^. Fusion of nuclease-like domains with short pAgos might compensate for the latter’s inactive PIWI endonuclease activity. Short pAgos might therefore have potential as new biotechnological tools^11^.

The *Crenotalea thermophila* short prokaryotic Ago/TIR-APAZ (SPARTA) system is such a short pAgo immune system that contains two components: a short pAgo protein and a TIR-APAZ protein containing a Toll/Interleukin-1 Receptor (TIR) domain fused to APAZ domain, which form a stable heterodimeric complex, known as SPARTA complex. Upon detection of invading DNA, the SPARTA complex oligomerizes and degrades essential NAD(P)^+^ to noncyclic ADPR(P) and NAM, leading to cell death^12^. The RNA-guided, sequence-specific DNA detection ability of the SPARTA complex enabled its repurposing as a programmable DNA detection tool^12^. Compared to CRISPR-Cas-based DNA detection tools, the SPARTA complex has a similar detection limit, does not require a protospacer adjacent motif in the target sequence, and enables detection of shorter genes, making it a promising new tool. However, the molecular mechanisms by which SPARTA recognizes specific target DNA and how target DNA binding activates SPARTA NADase activity remain unknown.

We used cryogenic electron microscopy (cryoEM) to determine the structures of the inactive monomeric *C. thermophila* SPARTA complex and the active tetrameric SPARTA complex bound to ssDNA targets. Supported by biochemical data, our structural analyses reveal mechanisms by which a small gRNA facilitates recognition of the complementary ssDNA target by the SPARTA complex and ssDNA target binding triggers SPARTA tetramerization and subsequent NADase activation. We also identify key residues responsible for the latter activities. Our results provide a mechanistic understanding of prokaryotic SPARTA-mediated immunity against invading plasmids.

## Results

### Target ssDNA binding triggers oligomerization of the SPARTA^gRNA^ complex

A short pAgo interacts with an associated TIR-APAZ protein to form a heterodimeric SPARTA complex, upon detection of invasive ssDNA, the SPARTA complex oligomerizes and subsequently degrades NAD(P)^+^^12^. To explain the molecular basis of these activities, we first assembled the SPARTA complex as indicated by previous studies^12^. We co-expressed the *C. thermophila* pAgo and TIR-APAZ proteins (Fig. 1a) in *Escherichia coli* and purified them from the cell lysate. Subsequent nickel affinity and gel filtration chromatography analyses revealed that pAgo and TIR-APAZ formed a stable heterodimeric complex, known as the monomeric apo SPARTA complex (Extended Data Fig. 1a). Size exclusion chromatography analysis showed that addition of gRNA and a complementary ssDNA target to the purified SPARTA complex *in vitro* resulted in an additional elution peak (Extended Data Fig. 1b). This peak did not appear when only gRNA was added (Extended Data Fig. 1b) and likely represents an oligomeric SPARTA complex. These results indicate that binding of target ssDNA stimulates SPARTA^gRNA^ oligomerization, consistent with previous observations^12^.

**Fig. 1.**
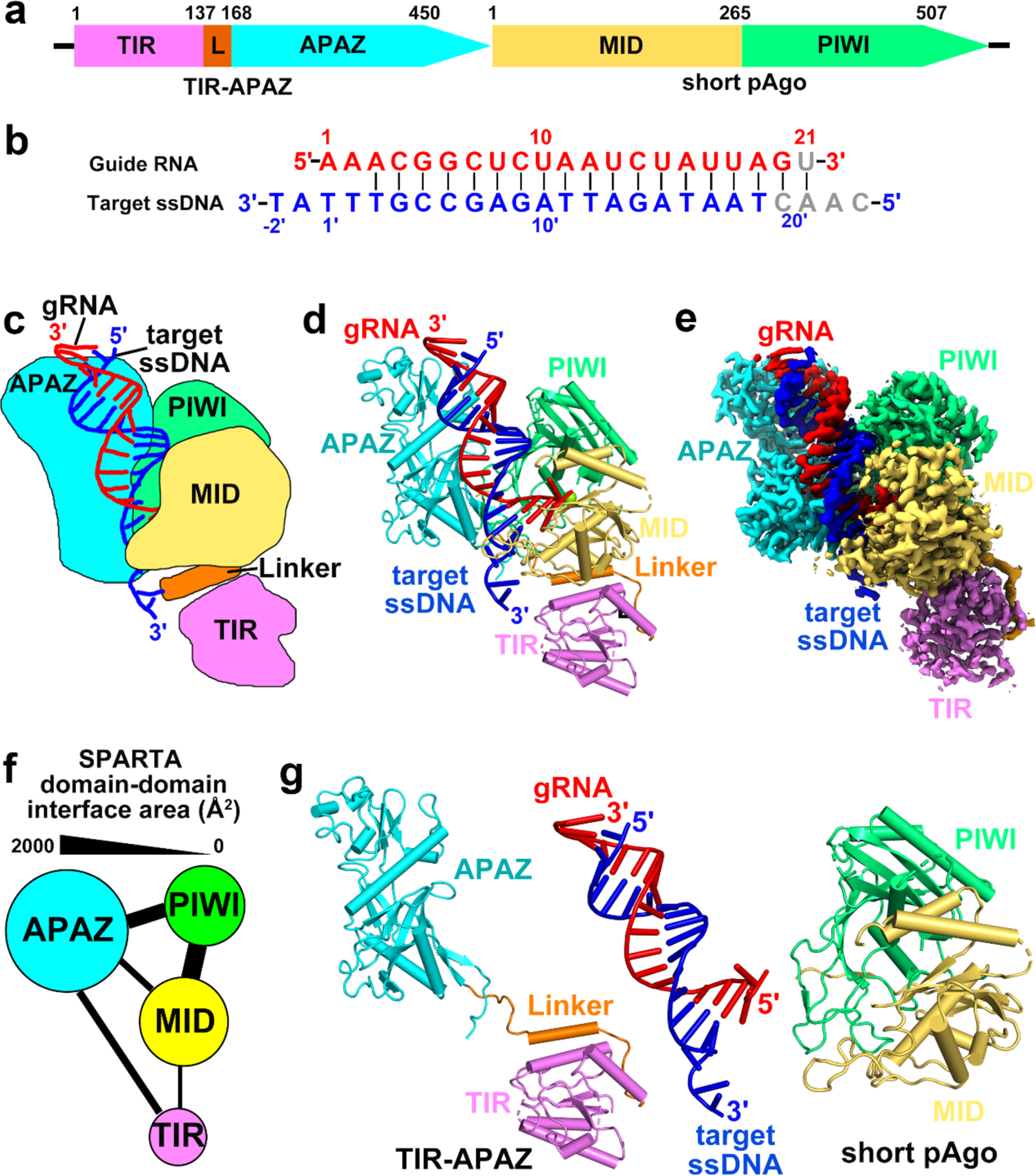
Overall architecture of monomeric *C. thermophila* SPARTA^gRNA^-target ssDNA complex. (a) The SPARTA system consists of TIR-APAZ and short pAgo proteins. (b) Schematic representation of pairing between the 5′-P gRNA (red) and target ssDNA (blue). Segments that can be traced are in color, while disordered segments are in grey. (c–e) Schematic (c), ribbon (d) and surface (e) representations of the 2.54-Å cryoEM structure of monomeric SPARTA^gRNA^-ssDNA target complex. (f) The domain-to-domain interaction network between TIR-APAZ and pAgo proteins in the SPARTA^gRNA^-ssDNA target complex. (g) Individual subunits of the SPARTA^gRNA^-ssDNA target complex. TIR-APAZ contains the APAZ and TIR domains, while the short pAgo contains the conserved MID and PIWI domains.

### The SPARTA complex contains a structurally conserved Ago architecture but a unique TIR effector domain

To validate the above results and investigate the molecular basis of SPARTA assembly and oligomerization, we performed cryoEM studies on the size exclusion chromatography elution sample we presumed to represent the oligomeric SPARTA^gRNA^-ssDNA complex (Extended Data Fig. 1b). Structural analysis resulted in classification of this sample into three different SPARTA^gRNA^-ssDNA complexes: a monomeric SPARTA^gRNA^-ssDNA complex at 2.54-Å resolution and two tetrameric SPARTA^gRNA^-ssDNA complexes at 3.13-Å and 2.92-Å resolution (Extended Data Fig. 2a).

**Fig. 2.**
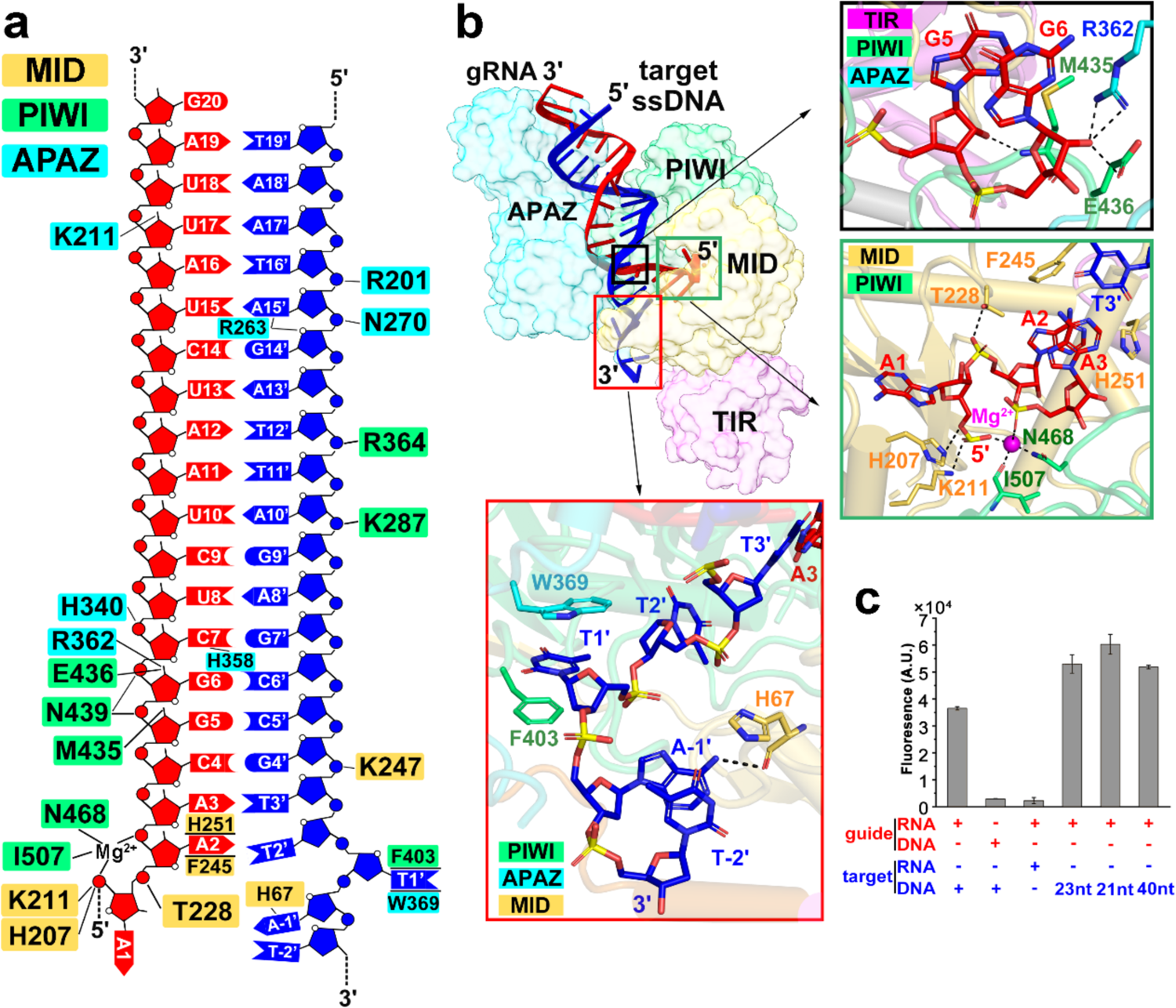
Recognition of the gRNA-ssDNA target duplex by the SPARTA complex. (a) Schematic interactions between SPARTA and gRNA (red) / ssDNA (blue) target duplex. Protein residues are colored according to domain. Hydrogen bonds are indicated by black lines, and the stacking interactions are shown by transverse lines. (b) Close-up view of SPARTA’s interactions with the 2′-OH of the gRNA (black inset), the 5′-P of the gRNA (green inset) and the 3′ end of the ssDNA target (red inset). (c) NADase activity of the SPARTA complex in the presence of the specified guide and target nucleic acids. This experiment was performed at least three times. Error bars represent standard deviation.

The higher-resolution, monomeric SPARTA^gRNA^-ssDNA complex adopts a bilobed architecture: one lobe contains the TIR domain connected to the APAZ domain by a short Linker, while the other lobe consists of a pAgo subunit that contains the conserved MID and PIWI domains (Fig. 1c–e, g and Extended Data Fig. 3). These two lobes forms an extensive interaction area with a buried interface of ∼2600 Å^2^ (Fig. 1f). The gRNA-target ssDNA duplex is bound in the positively charged channel between these two lobes (Fig. 1c–e). We observed a 20-nucleotide traceable segment of the 21-nucleotide gRNA and a 21-nucleotide traceable segment of a 25-nucleotide complementary ssDNA target in the gRNA-ssDNA heteroduplex (Fig. 1b), which formed extensive contacts with all domains except the TIR domain (Fig. 1c–e). Structural comparison via distance matrix alignment revealed that the monomeric SPARTA^gRNA^-ssDNA complex structurally resembles the long prokaryotic *Rhodobacter sphaeroides* Ago (RsAgo) protein (PDB 5AWH) involved in RNA-guided DNA silencing^13^, with a Cα root mean square distance (RMSD) value of 5.1 Å (Extended Data Fig. 4a). Structure and sequence alignments revealed that SPARTA lacks the conserved catalytic tetrad motif (DEDX) within the PIWI domain that is responsible for target nucleic acid cleavage, suggesting that SPARTA does not catalyze target ssDNA cleavage^5, 14^ (Extended Data Fig. 4b, c). Previous biochemical studies on the SPARTA system have also demonstrated the absence of its target ssDNA cleavage activity^12^. Consistent with this finding, our structure of the monomeric SPARTA complex revealed a consecutive density of target ssDNA without being cleaved in Extended Data Fig. 3. Thus, our determined SPARTA^gRNA^-ssDNA complex further demonstrated that monomeric SPARTA lacks the ability to cleave target ssDNA and is therefore inactive.

**Fig. 3.**
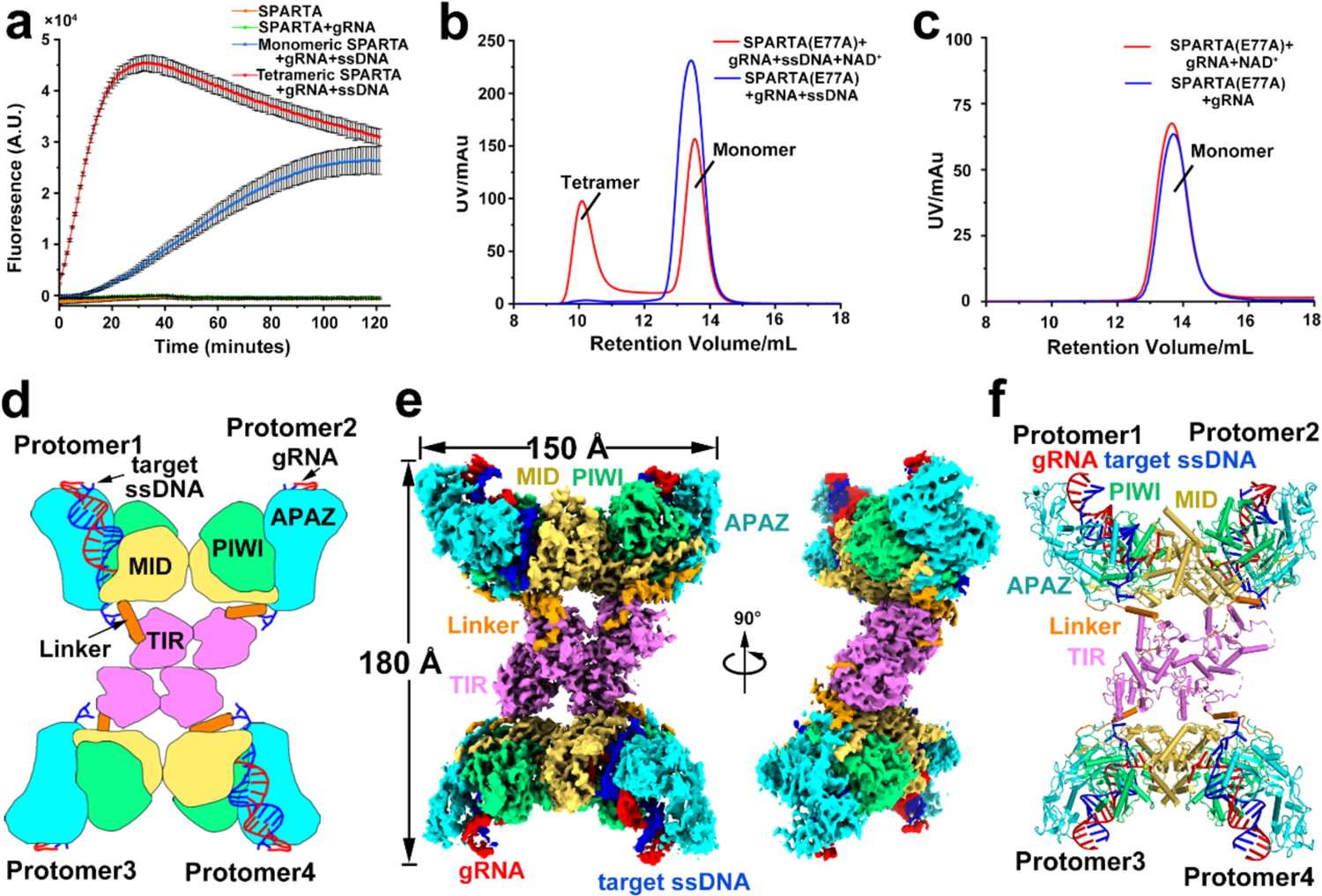
Overall architecture of the tetrameric SPARTA^gRNA^-ssDNA complex. (a) NADase activity of SPARTA, SPARTA^gRNA^ and monomeric and tetrameric SPARTA^gRNA^-ssDNA was monitored over time. (b, c) Size exclusion chromatography profiles of the catalytically inactive SPARTA^gRNA^ E77A mutant incubated with 5 mM NAD^+^ in the presence (b) and absence (c) of target ssDNA. (d–f) Schematic (d), surface (e) and ribbon (f) representations of the tetrameric SPARTA^gRNA^-ssDNA complex.

**Fig. 4.**
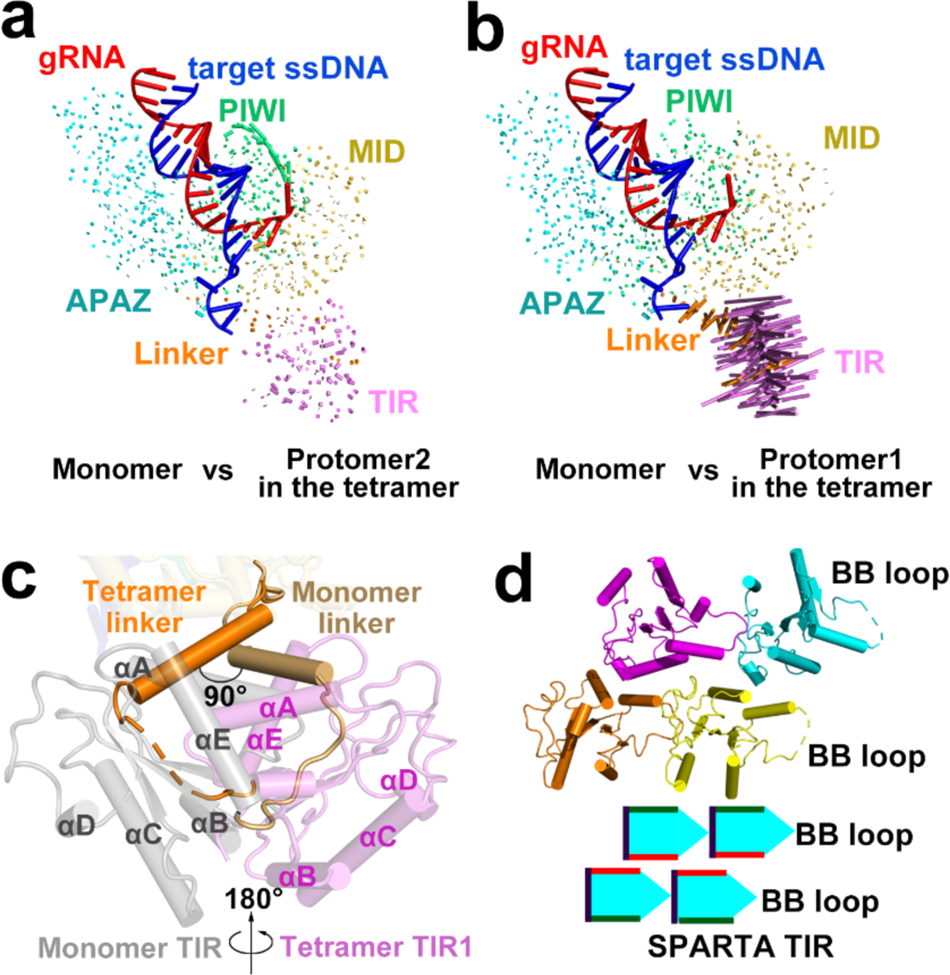
Target ssDNA binding induces conformational changes in TIR domains. (a, b) Structural comparison of the monomeric SPARTA complex with the tetrameric SPARTA complex protomer 2 (a) and protomer 1 (b). Vector length correlates with the domain movement scale. (c) The Linker and TIR domains undergo conformational rearrangement upon SPARTA tetramer complex formation: the Linker domain undergoes a 90° rotation and the TIR domain undergoes a 180° rotation. (d) Ribbon and schematic representations of the TIR domain assembly pattern in tetrameric SPARTA complex.

Distinct from previously reported Ago proteins, the SPARTA complex contained an extra TIR domain fused to APAZ (Extended Data Fig. 4a), which structurally resembled the N domain rather than the PAZ domain of RsAgo (Extended Data Fig. 4d). Structural comparison revealed that the TIR domain of SPARTA resembled the myeloid differentiation primary response gene 88 (MyD88) TIR domain with an RMSD of 3.4 Å^15^ (Extended Data Fig. 4e). The TIR domain had a conserved Rossman-like fold pattern, which consists of five-stranded parallel β-sheets (βA to βE) typically with five surrounding α-helices (αA to αE) separated by loops^16, 17^ (Extended Data Fig. 4f). The functionality of TIR domains typically requires the formation of the high-order oligomers, which suggests that the TIR domain of SPARTA might oligomerize for NAD^+^ hydrolysis^12, 18^.

### Structural basis of SPARTA’s gRNA-ssDNA target heteroduplex binding preference

SPARTA degrades NAD^+^ more efficiently when it binds to gRNA containing a 5′-phosphate (5′-P) group and its complementary ssDNA than when it forms other combinations (5′-hydroxyl (5′-OH) gRNA-ssDNA target duplex, 5′-P guide DNA (gDNA)-single-stranded RNA (ssRNA) target duplex, or 5′-OH gDNA-ssRNA target duplex)^12^. This finding suggests that SPARTA preferentially uses a small RNA containing a 5′-P group as a guide to recognize its ssDNA target. Examining our structure of the SPARTA^gRNA^-ssDNA complex, we observed that SPARTA recognized the gRNA-target ssDNA heteroduplex mainly through contacts with the sugar-phosphate backbones and to a smaller extent via stacking interactions with nucleotide bases (Fig. 2a). Koopal.et al previously showed that the SPARTA exhibits a preference for binding small RNAs with a 5’-AA dinucleotide^12^. In our determined SPARTA complex, we observed that the residues F245 and H251 within the MID domain of SPARTA stack with the second adenine of the gRNA strand (Fig. 2a, b, green inset). However, we were unable to identify the specific residue responsible for the preference for the first adenine of the gRNA strand, likely due to the intrinsic flexibility of the region spanning residues 146-203 within the MID domain. Moreover, the 2′-OH groups of the nucleotides at positions 5, 6 and 17 in the gRNA were specifically recognized by residues M435^PIWI^, E436^PIWI^, R362^APAZ^ and K211^APAZ^ through hydrogen-bonding interactions (Fig. 2a, b, black inset). These findings highlight a lack of sequence specificity for SPARTA’s recognition of the guide-target duplex. The 5′-P group of the gRNA was stabilized by residues H207 and K211 as well as by a divalent cation in the MID domain, which neutralized the negative charge of the 5′-P (Fig. 2b, green inset). This specific 5′-P recognition mode explains the reported binding preference of the SPARTA complex for gRNA with a 5′-P rather than a 5′-OH group^12^. Consistent with this structural analysis, replacement of gRNA with gDNA markedly reduced SPARTA-mediated NADase activity (Fig. 2c), demonstrating that SPARTA preferred to bind RNA as its nucleic acid guide and explaining why SPARTA prefers transcriptome-derived gRNAs with a 5′-P *in vivo*^12^.

At the 3′ end of the target ssDNA, nucleotides T2′ to T−2′ are unpaired and splayed outwards. The splayed bases from position 2′ to −1′ were stabilized by hydrophobic stacking interactions (H67^MID^, F403^PIWI^ and W369^APAZ^) in the MID, PIWI and APAZ domains (Fig. 2a, b, red inset). Extending the 3′ end of the target ssDNA to 40 nucleotides, or shortening it by deleting nucleotides T−2′ to A−1′ (23 nucleotides) or T−2′ to T2′ (21 nucleotides), had negligible effect on SPARTA NADase activity (Fig. 2c). However, substitution of target ssDNA with target ssRNA dramatically impaired SPARTA NADase activity (Fig. 2c), indicating that ssDNA is the preferred target nucleic acid for the SPARTA^gRNA^ complex. However, we did not observe any specific contacts with the 2′-groups of the target ssDNA and the protein residues, which prevent us from providing evidence for the preference of target ssDNA over target ssRNA. Taken together, our results demonstrate that SPARTA prefers to bind a gRNA-target ssDNA heteroduplex through sequence-nonspecific contacts, with specific recognition of the 5′-P and 2′-OH groups of the gRNA. Our findings provide the structural evidence for a previous observation that SPARTA prefers to recruit a small gRNA with a 5’-P group *in vivo*^12^.

### Target ssDNA-induced tetramerization activates SPARTA NADase activity

The oligomerization of *Maribacter polysiphoniae* SPARTA is required for its NADase activity^12^. To determine whether for the same is true for *C. thermophila* SPARTA, we evaluated NADase activity of the SPARTA^gRNA^-ssDNA complex in its monomeric and tetrameric states via an *in vitro* fluorescence-based assay. Removal of the nicotinamide group of ε-NAD^+^ (a fluorescent NAD^+^ analog) leads to a pronounced fluorescence signal, which can be used to measure the NADase activity. We incubated ε-NAD^+^ with the individual, purified SPARTA complexes and monitored the change in fluorescence conferred by SPARTA-mediated ε-NAD^+^ consumption (Fig. 3a). The tetrameric SPARTA^gRNA^-ssDNA complex demonstrated pronounced and immediate NADase activity following addition of ε-NAD^+^, while the monomeric SPARTA^gRNA^-ssDNA complex consumed ε-NAD^+^ at a slower rate (Fig. 3a). No NADase activity was observed for the apo SPARTA complex regardless of the addition of gRNA, indicating that target ssDNA is an essential trigger for SPARTA NADase activity (Fig. 3a).

Given that self-association of the TIR domain is typically essential for NADase activity^19^. we speculated that the observed weak NADase activity of the monomeric SPARTA^gRNA^-ssDNA complex might be due to oligomerization of the SPARTA^gRNA^-ssDNA complex upon addition of NAD^+^. To test this hypothesis, we incubated 5 mM NAD^+^ with the monomeric SPARTA^gRNA^-ssDNA complex containing an E77A mutation in the TIR domain, which cannot perform NAD^+^ hydrolysis^12^. Subsequent size exclusion chromatography analysis revealed that a significant portion of the SPARTA^gRNA^-ssDNA monomeric complex had converted into the tetrameric form (Fig. 3b), demonstrating that NAD^+^ binding stimulates tetramerization of the SPARTA^gRNA^-ssDNA complex. In contrast, the addition of NAD^+^ to the monomeric SPARTA complex in the absence of target ssDNA did not result in tetramerization of the SPARTA^gRNA^ complex (Fig. 3c), indicating that target ssDNA binding is essential for SPARTA tetramerization. Collectively, our biochemical data show that target ssDNA binding is required for SPARTA tetramerization, which is further promoted by the presence of NAD^+^ substrate and enables NAD^+^ hydrolysis.

### The activated SPARTA^gRNA^-ssDNA complex assembles into a tetrameric state

CryoEM analysis of the oligomeric SPARTA eluate from the gel filtration chromatography analysis yielded reconstruction of the tetrameric SPARTA^gRNA^-ssDNA complex in two different assembly states with overall resolutions of 3.13 Å and 2.92 Å (Fig. 3d–f and Extended Data Fig. 2). To build an atomic model of the tetrameric SPARTA^gRNA^-ssDNA complex, we initially docked four copies of the monomeric SPARTA^gRNA^-ssDNA complex model into the cryoEM density maps of two reconstructed tetrameric complexes by rigid body fitting. The resulting model was then further refined according to the density maps. Upon aligning the resulting structures of this different assembly states, we observed that the SPARTA tetramer in state 2 aligned well with the SPARTA tetramer in state 1 through a 180° rotation, probably due to the alignment differences in different dimers of the SPARTA tetramer during the cryoEM data processing (Extended Data Fig. 5a). We then chose the structure of tetrameric SPARTA^gRNA^-ssDNA complexes in state 1 for further analysis. The tetrameric SPARTA^gRNA^-ssDNA complexes contains four asymmetric copies of the SPARTA^gRNA^-ssDNA complex protomer. The overall architecture of the tetrameric SPARTA complex is arranged in a “dimers of dimers” X-shape, whereby protomers 1 and 2, as well as protomers 3 and 4, formed basic dimers through short pAgo-mediated interactions in a buried interface of ∼1300 Å^2^. Two such dimers were then arranged into a tetrameric state via different interactions between the two TIR-interfacing dimers with a buried interface of ∼1080 Å^2^. The four TIR domains formed an asymmetric, two-stranded, head-to-tail, parallel arrangement at the center of the complex, while the remaining domains, along with the gRNA-ssDNA target duplex, were positioned outside the TIR complex (Fig. 3d–f).

**Fig. 5.**
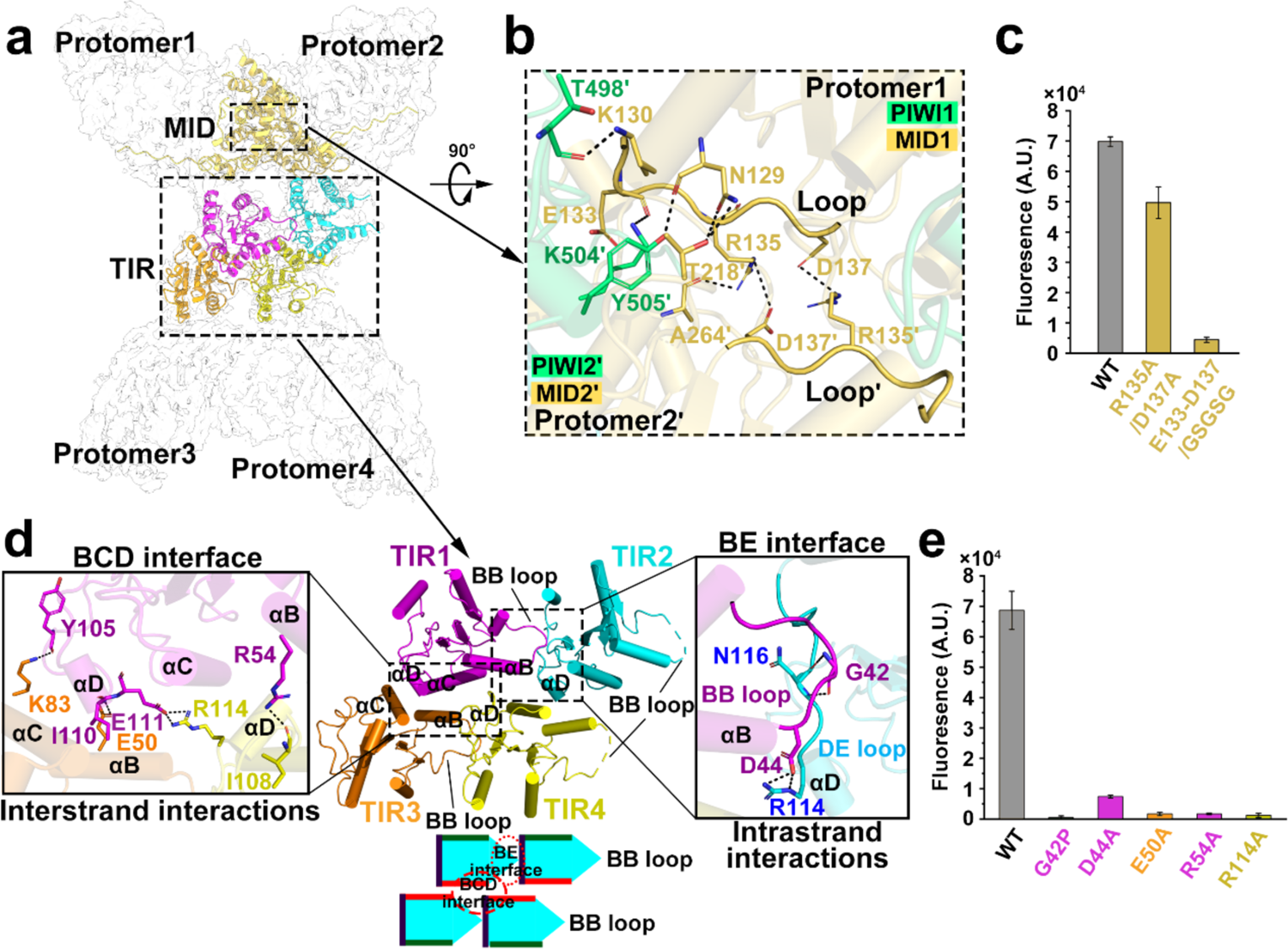
Formation of a SPARTA tetramer is mediated by pAgo and TIR domain interactions. (a) A global view of the SPARTA tetramer. Cartoon representation of two MID domains and four TIR domains, colored. (b) Detailed interactions between the two pAgos from protomer 1 and protomer 2 of the tetrameric SPARTA complex. Hydrogen bonds are indicated by black lines, and the protein residues are colored according to domain. (c, e) Effect of pAgo (c) and TIR (e) interface mutants on NADase activity. (d) Detailed interactions between the four TIR domains of the tetrameric SPARTA complex at the BCD (left panel) and BE (right panel) interfaces. Hydrogen bonds are indicated by black lines, and the protein residues are colored according to domain.

### Conformational changes in the TIR domains of the SPARTA^gRNA^-ssDNA complex are required for NADase activation

Tetramerization of the SPARTA complex is essential for its NADase activity, the monomeric SPARTA complex is inactive (Fig. 3a, b)^12^. Comparison of the structure of the inactive monomeric SPARTA^gRNA^-ssDNA complex to each protomer in the active tetrameric SPARTA^gRNA^-ssDNA complexes revealed that the four protomers are not identical and adopt two different conformations. Compared to the inactive monomeric SPARTA complex, protomers 2 and 4 adopt an identical conformation with an RMSD of 0.6 Å (Fig. 4a), while the Linker and TIR domains in protomers 1 and 3 undergo 90° and 180° rotations, respectively, enabling tetramerization of the TIR domains (Fig. 4b, c). Rearrangement of the TIR domains in protomers 1 and 3 enables the four TIR domains to assemble into two parallel, head-to-tail dimers, which interact with each other (Fig. 4d). This TIR arrangement was reminiscent of the TIR assembly pattern in MyD88 (Extended Data Fig. 5b), a eukaryotic scaffold protein involved in immune signaling that does not harbor NADase activity^20^. However, the SPARTA TIR arrangement was different from that of the eukaryotic SARM1 enzyme (Extended Data Fig. 5b), even though the latter possesses NADase activity^19, 21^. The recently reported prokaryotic protein *Acinetobacter baumannii* TIR (AbTIR) possesses the eukaryotic signaling scaffold assembly but is NADase active (Extended Data Fig. 5b)^22^. These analyses suggest that the eukaryotic classification of the scaffold and enzyme assemblies for TIR domains may not be appliable to the prokaryotic TIR domains.

### Formation of the NAD^+^ catalytic pocket in the SPARTA TIR tetramer assembly

Upon RNA-guided binding of target ssDNA, SPARTA tetramerized through interactions mediated by both the short pAgo and the TIR domain, causing a disordered loop region (N129 to D137) in the MID domain of short pAgo to become ordered via interactions with the adjacent short pAgo (Fig. 5a, b and Extended Data Fig. 6a). This loop was much longer than its equivalent in other reported monomeric Ago proteins (Extended Data Fig. 6b, c)^23^, suggesting that it might play a role in facilitating pAgo dimerization. This short pAgo loop consists of hydrophilic residues with long side chains (129-NKNDEERVD-137) that interact with the MID and PIWI domains of the adjacent pAgo subunit through salt bridges (R135^MID^ and D137^MID^) and hydrogen bonds (T218^MID^, A264^MID^, T498^PIWI^, K504^PIWI^, Y505^PIWI^) (Fig. 5b). Mutation of residues R135 and D137 to alanine, or replacement of this loop (133-EERVD-137) with a GSGSG linker, severely impaired SPARTA NADase activity (Fig. 5c). These findings demonstrate that pAgo loop-mediated interactions play a crucial role in promoting SPARTA NADase activity.

**Fig. 6.**
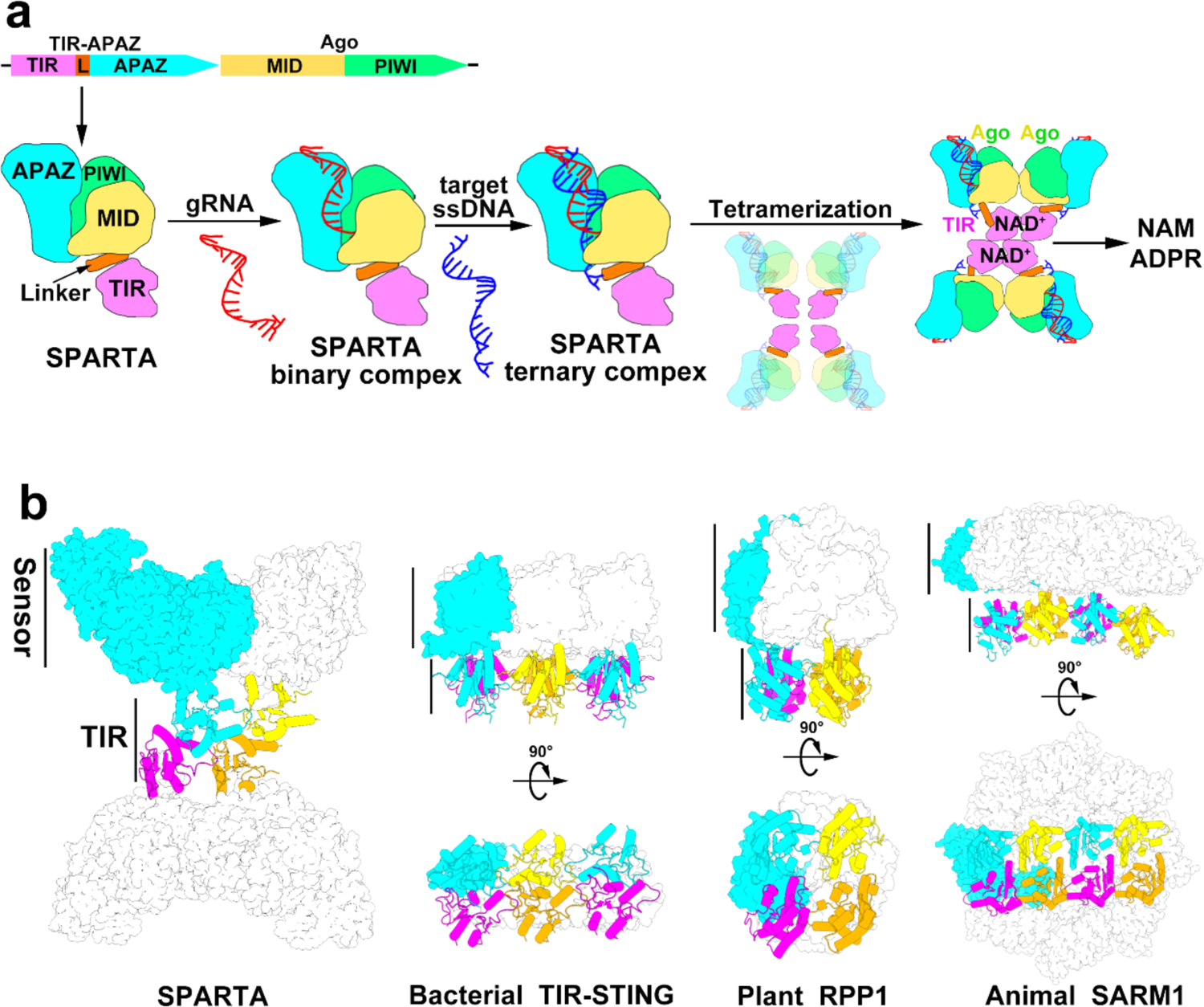
Proposed mechanistic model of prokaryotic SPARTA complex activation in response to invading plasmid DNA. (a) Proposed mechanistic model for oligomerization-dependent SPARTA activation. The short Ago and TIR-APAZ proteins, encoded by genes in the same operon, assemble into the SPARTA complex. The SPARTA complex binds a small 5′-P gRNA, which enables recognition of complementary target ssDNA. Target ssDNA binding causes conformational changes in a specific loop of the MID and TIR domains, resulting in the formation of a SPARTA tetramer complex via pAgo- and TIR-mediated interactions. Tetramerization of the TIR domains creates two NAD^+^ catalytic pockets that are primed for the hydrolysis of NAD^+^ into NAM and ADPR. (b) Structural comparison of TIR domain-containing proteins in their active states, including bacterial SPARTA, bacterial *sf*TIR-STING (PDB 7UN8), plant RPP1 (PDB 7DFV) and animal SARM1 (PDB 7NAK, TIR domains; PDB 7NAL, sensor domains). TIR domains are represented in different colors, and one protomer of each sensor is highlighted in cyan.

The aforementioned loop-mediated interactions in SPARTA caused dimerization of the short pAgo, bringing the associated TIR domains into proximity to form a basic, asymmetric, head-to-tail TIR dimer via interactions between the BB and DE loops (Fig. 5d, right inset), which comprised a buried BE interface of ∼480 Å^2^. Interactions with the adjacent DE loop caused the BB loop to become ordered in TIR1 and TIR3, whereas it remained disordered in TIR2 and TIR4 (Fig. 5d). Rearrangement of the BB loop at the BE interface facilitates NAD^+^ access to its binding and catalytic sites and subsequent NAD^+^ degradation^24–27^. G42P or D44A mutations in the BE interface almost completely abolished NAD^+^ hydrolysis (Fig. 5e), confirming the essential role of the BE interface in TIR NADase activity.

NAD^+^ binding sites span two TIR domains in AbTIR, with W227^TIR^^2^ stacking the adenine group, and the conserved catalytic residues E208^TIR^^1^ and W204^TIR1^ promoting NAD^+^ hydrolysis and cyclization of the ADPR product, respectively^22^. We wondered whether NAD^+^ binding sites also span two TIR domains in *C. thermophila* SPARTA. To investigate this possibility, we docked the NAD^+^ mimic 3AD (8-aminoisoquinoline adenine dinucleotide) into the catalytic pocket on the BE surface by superimposing the TIR1 and TIR2 domains of SPARTA with AbTIR^22^ (Extended Data Fig. 6d). The aromatic residue Y105^TIR2^ stacked the adenine group, while the conserved catalytic residue E77^TIR1^ was primed to facilitate NAD^+^ hydrolysis (Extended Data Fig. 6e). The SPARTA TIR lacks the aromatic residue essential for ADPR cyclization (W204 in AbTIR)^22^, and instead contains a glycine in the corresponding position (G73). This observation supports the previous finding that the SPARTA complex hydrolyzes NAD^+^ into ADPR rather than cyclic ADPR.

The TIR intra-strand BE interface forms the NAD^+^ binding sites, while the interstrand BCD interface connects the two parallel, head-to-tail strands together via the TIR αB, αC and αD helices. Mutating specific residues (E50, R54 and R114) in the BCD interface to alanine eliminated NADase activity (Fig. 5e), confirming the essential role of the BCD interface in TIR NADase activation. Size exclusion chromatography analysis of these mutants showed that they assembled in a state similar to the wild type SPARTA (Extended Data Fig. 7), indicating that the loss of the NADase activity in these mutants was not due to misfolding. Generally, the BE and BCD interfaces of TIR domains form filament structures^19^. However, the unique pAgo-mediated interactions within the SPARTA complex prevented its TIR domains from forming an open-ended filament, allowing only an assembly state (Fig. 3f, i).

Collectively, our data show that tetramerization of SPARTA is facilitated by the BE and BCD interfaces of the TIR domain as well as by the pAgo dimer interface, and all of these interfaces are essential for NAD^+^ hydrolysis. The intact catalytic pocket necessary for NAD^+^ hydrolysis is formed by bridging of the two adjacent TIR domains via tetramerization. These findings are consistent with our observation that the addition of NAD^+^ promotes SPARTA tetramerization (Fig. 3b).

## Discussion

Ago proteins use small oligonucleotides as guides to bind complementary DNA/RNA, mediating eukaryotic gene silencing and prokaryotic defense against invading nucleic acids^1,2^. Other proteins contain TIR domains that serve as either signaling scaffolds or NADase enzymes involved in innate immunity and cell death across all domains of life^19^. The recently identified prokaryotic SPARTA immune systems link TIR domains with Ago proteins. Detection of invading plasmid DNA by the SPARTA pAgo activates its associated TIR domain, enabling NADase activity. Thus, the SPARTA system has great potential as a DNA detection tool^12^. However, the molecular details underlying target ssDNA-mediated activation of TIR NADase activity have been elusive. In this study, we determined the cryoEM structures of inactive monomeric and active tetrameric *C. thermophila* SPARTA complexes. Supported by biochemical data, we propose a mechanistic model for the SPARTA-mediated immune response to invading DNA (Fig. 6a).

The prokaryotic SPARTA immune system relies on two components, a TIR-APAZ protein and a short pAgo, to achieve SPARTA-mediated immunity through TIR domain-mediated depletion of NAD^+^ in response to invasive DNA (Fig. 6a)^12^. The short pAgo consists of MID and PIWI domains, which directly interact with the APAZ domain in TIR-APAZ, and forms a heterodimeric SPARTA complex (referred to as monomeric apo SPARTA) that cannot perform NAD^+^ hydrolysis. The monomeric SPARTA complex prefers to recruit small RNAs with a 5′-P group as guides *in vivo*^12^. This preference is due to specific recognition of the 5′-P and 2′-OH groups of gRNA by the MID, PIWI and APZA domains (Fig. 2a, b).

Upon binding to the complementary ssDNA target, the SPARTA complex undergoes significant conformational changes: short pAgo dimerizes, bringing the associated TIR domains together to form two basic TIR dimers (Fig. 5a). Within each TIR dimer, one TIR domain undergoes a 180° rotation (Fig. 4c), leading to rearrangement of the BB loop and generation of the NAD^+^ catalytic pocket, which spans two adjacent TIR domains (Extended Data Fig. 5d). The two pairs of TIR domains then self-associate via the BCD interface to form a two-stranded assembly pattern (Fig. 5d). The BE and BCD interfaces are essential for NADase activity, as mutation of specific residues of either interface greatly impair SPARTA NADase activity (Fig. 5e). Upon TIR tetramerization, two NAD^+^ catalytic pockets are formed. The SPARTA TIR domains lack an essential aromatic residue (W204 in AbTIR) that mediates ADPR cyclization and instead contain a glycine residue (G73) in the corresponding position (Extended Data Fig. 6e). Thus, NAD^+^ hydrolysis by SPARTA produces ADPR and NAM. Degradation of essential NAD^+^ triggers suicide of the invaded cell, thereby protecting the rest of the bacterial population^12^.

The TIR domain plays an important role in the innate immune systems of all three domains of life. Its primary function is an effector that transmits an immune signal upon detection of invasive signals via its associated sensor domain. Typically, the functionality of TIR domains requires their self-association, but due to the weak affinity between TIRs, their association often requires assistance from sensor domains^18^. Self-association of SPARTA TIRs is facilitated by the short pAgo, which serves as a sensor for invading DNA (Fig. 6b). The TIR domain functions as an effector by depleting NAD^+^ to trigger bacterial abortive infection^12^. Prior to signal detection by the short pAgo, TIR NADase activity is repressed by keeping the TIR domains apart. Thus, in prokaryotic SPARTA immune systems, TIR oligomerization-dependent NADase activity is tightly regulated by its associated sensor domain.

TIR domains serve as an essential component of other prokaryotic immune systems, including cyclic nucleotide-based antiphage signaling and Thoeris and Pyscsar (pyrimidine cyclase system for antiphage resistance) defense systems^22, 25, 28–32^. The immune response of these systems also requires a sensor domain to activate the TIR effector domains and degrade essential NAD^+^. For example, binding of the signal cyclic di-GMP by the STING sensor domain drives the formation of oligomeric, open-ended TIR filaments, rearranging the NAD^+^ catalytic site to initiate NAD^+^ hydrolysis and abortive infection^29^ (Fig. 6b). In plants, the TIR domain of RPP1 tetramerizes and degrades NAD^+^ upon recognition of the pathogen-derived effector ATR1 by the C-terminal jelly roll/Ig-like sensor domain, ultimately leading to cell death^27^ (Fig. 6b). In animals, the ARM domain in SARM1 detects the allosteric activator NMN and triggers the self-association of TIR domains^21^ (Fig. 6b).

Although TIR domain-containing proteins possess a conserved glutamic acid residue necessary for TIR-mediated NAD^+^ cleavage, the reaction products vary among different TIR-containing systems. For instance, while SARM1 produces either ADPR or cADPR, bacterial and plant TIR domains degrade NAD^+^ into ADPR and various isomers of cADPR, indicating that the catalytic mechanism varies in different species^18^. SPARTA TIR domains lack the aromatic residue essential for ADPR cyclization, leading to the hydrolysis of NAD^+^ into ADPR rather than cADPR. cADPR typically functions as a second messenger to activate the downstream immune signaling pathway^22^. Thus, TIR domains can participate in the immune signaling pathway differently, either by depleting the essential metabolite NAD^+^ or by generating new signaling molecules (e.g., cADPR).

In conclusion, our study elucidates the molecular underpinnings of prokaryotic SPARTA immune systems that respond to invading plasmid DNA. The short pAgo and TIR-APAZ proteins form a stable SPARTA effector complex. The short pAgo serves as a sensor to bind a small RNA guide, which recognizes its complementary ssDNA target. Binding of target ssDNA by the short pAgo triggers SPARTA tetramerization and rearrangement of its TIR domains, leading to the formation of a catalytic pocket where NAD^+^ hydrolysis subsequently occurs. These findings deepen our understanding of prokaryotic SPARTA defense systems and provide a molecular basis for development of SPARTA-based biotechnological tools.

## Methods

### Bacterial strains

*Escherichia coli* DH5α and BL21 Star (DE3) cells were used for plasmid reconstruction and protein expression, respectively.

### Protein expression and purification

The plasmid encoding Crenotalea thermophila SPARTA was a gift from Daan Swarts (Addgene plasmid #183146; http://n2t.net/addgene:183146; RRID: Addgene_183146). SPARTA vector was transformed into *Escherichia coli* BL21 Star (DE3), a single colony was picked to inoculate a 200 mL LB culture, which was grown at 200 rpm and 37 °C. This culture was further used to inoculate 12×1000 mL LB medium which was incubated at 200 rpm and 37 °C until OD600 of ∼ 0.6 was reached. The SPARTA proteins expression was induced by adding isopropyl-β-D-1-thiogalactopyranoside to a final concentration of 0.5 mM at 18 °C for 20 hrs. Cells were harvested by centrifugation at 5,000 ×g for 10 min and resuspended in lysis buffer (20 mM Tris-HCl pH 7.5, 500 mM NaCl, 20 mM imidazole, 1 mM DTT). The harvested cells were lysed by sonication (Avestin EmulsiFlex-C3 homogenizer) and centrifuged for 30 min at 16,000 rpm. The supernatant was applied to a 5 mL HisTrap Fast flow column (Cytiva life Sciences). The column was then washed with 10 column volumes (CV) of lysis buffer and protein was eluted with elution buffer (20 mM Tris-HCl pH 7.5, 500 mM NaCl, 300 mM imidazole, 1 mM DTT). The elution fractions were pooled and TEV protease was added in a 1:50 ratio, then incubated at 16 °C overnight to remove the MBP tag. The proteins were diluted with 20 mM Tris-HCl pH 7.5 and applied on a 5 mL HiTrap Heparin column (Cytiva Life Sciences). The proteins were eluted by a linear gradient from 100 mM to 1 M NaCl in 20 CVs, and then concentrated using 50 kDa molecular mass cut-off concentrators (Amicon) before further purification over a Superdex 200 increase 10/300 GL column (Cytiva Life Sciences) pre-equilibrated in size-exclusion chromatography (SEC) buffer (20 mM Tris-HCl pH 7.5, 150 mM NaCl, 2 mM MgCl_2_, 1 mM DTT). Peak fractions were assessed by gel electrophoresis, and were pooled and concentrated, flash frozen in liquid nitrogen and stored at −80 °C.

All mutants were generated by site-directed mutagenesis and purified by the same methods as mentioned above.

### *In vitro* assembly of SPARTA^gRNA^ and SPARTA^gRNA^-ssDNA target complexes

To assemble SPARTA^gRNA^ and SPARTA^gRNA^-ssDNA target complexes, the purified SPARTA complex was diluted to 1 mg/mL and mixed with 5’-phosphorylated guide RNA (5’P-AAACGGCUCUAAUCUAUUAGU-3’) at a molar ration of 1:1.2. After incubated at 55 °C for 15 min, DNA target (5’-CAACTAATAGATTAGAGCCGTTTAT-3’) was added at a molar ration of 1:1.5 (protein: DNA) and incubated at 55 °C for another 15 min. The sample was then loaded on a Superdex 200 Increase 10/300 GL column (Cytiva Life Sciences) which was pre-equilibrated in SEC buffer. Peak fractions were assessed by gel electrophoresis, and were pooled and concentrated, flash frozen in liquid nitrogen and stored at −80 °C.

### Cryo-EM sample preparation and data acquisition

A 3.5 μL concentrated SPARTA sample was loaded onto a glow discharged UltrAuFoil 300 mesh R1.2/1.3 grids (Quantifoil) using a FEI Vitrobot at 4°C and 100% humidity. The grids were transferred to FEI Titan Krios electron microscope operated at 300 kV, and movies (32 frames, total accumulated dose 50 e^-^/Å^2^) were collected using a direct electron detector Gatan K3 in the counting mode with a defocus range from −1.5 to −2.5 μm. Automated single-particle data acquisition was performed with SerialEM^33^ program in a nominal magnification of 105,000, yielding a final pixel size of 1.1 Å.

### Cryo-EM data processing

Image processing were performed by RELION 3.1^34^ and cryoSPARC v3.1^35^. The motion correction was performed with MotionCor2^36^. Contrast transfer function (CTF) parameters were estimated by CtfFind4^37^. A total of 2,571,055 particles were auto-picked using Laplacian-of-Gaussian and extracted using RELION 3.1. After several iterations of 2D classifications, particles from the good classes were selected and subjected to 3D reconstruction with the initial model generated by cryoSPARC v3.1 as a reference. Particles corresponding to the best class were selected and subjected to non-uniform refinement in cryoSPARC v3.1. All the cryo-EM reconstructions were estimated with the gold standard Fourier shell correlation using the 0.143 threshold^38^. Local resolution estimates were calculated from two half data maps in cryoSPARC v3.1. The details related to data processing were shown in Extended Data Table 1.

### Model building and refinement

With the assistance of bulky residues and PSIPRED secondary structure prediction^39^ together with structural models predicted by Robetta (https://robetta.bakerlab.org/), we manually built the atomic models interactively in COOT^40^. The real-space refinement in PHENIX^41^ was used to refine all models against the cryo-EM maps by applying geometric and secondary structure restraints. All structure Fig.s were prepared in PyMol (http://www.pymol.org) and ChimeraX^42^.

### *In vitro* NADase activity assays

Briefly, a reaction mixture contains the purified SPARTA complex (removal of MBP tag for Fig. 2c and 3a) or variants (with MBP tag for Fig. 5c and 5e) in SEC buffer, fluorescent NAD^+^ analog ε-NAD^+^ (Sigma, N2630), RNA guide, DNA target or variants, and 5×reaction buffer (50 mM MES pH 6.5, 375 mM KCl, and 10 mM MgCl_2_). The final concentrations of each component were 1 μM SPARTA complex, 50 μM ε-NAD^+^, 1.2 μM RNA guide, 1.2 μM DNA target, 10 mM MES pH 6.5, 125 mM KCl, and 2 mM MgCl_2_ in a final volume of 40 μL. Due to the temperature limitation of the instrument, reaction mixtures were took place at 45 °C (Fig. 3a) or incubated at 55 °C (Fig. 2c, 5c and 5e) for 1 hrs before added in 96-well half area plates (Corning, 3690). Fluorescence intensity was measured using an excitation wavelength of 300 nm and emission wavelength of 410 nm of each well in a Synergy H1 microplate reader (BioTek). All experiments were repeated at least three times and error bars indicate standard deviations.

### Data availability

The cryo-EM density maps have been deposited in the Electron Microscopy Data Bank (EMDB) under accession number EMD-35419 (monomeric SPARTA^gRNA^-ssDNA target complex); EMD-35420 (tetrameric SPARTA^gRNA^-ssDNA target complex in state1) and EMD-35421 (tetrameric SPARTA^gRNA^-ssDNA target complex in state2). The atomic coordinates have been deposited in the Protein Data Bank (PDB) with accession number 8IFK (monomeric SPARTA^gRNA^-ssDNA target complex); 8IFL (tetrameric SPARTA^gRNA^-ssDNA target complex in state1) and 8IFM (tetrameric SPARTA^gRNA^-ssDNA target complex in state2). This paper does not report original code. Any additional information required to reanalyze the data reported in this paper is available from the lead contact upon request.

## Acknowledgements

We thank the staff at Southern University of Science and Technology (SUSTech) Cryo-EM Center for assistance in data collection on the SUSTech Titan KRIOS cryo-electron microscope. This work was supported by the National Natural Science Foundation of China (Grant No. 32270050 to N.J.), Shenzhen and Guangdong Natural Science Foundation (Grant No. JCYJ20220530114409022 and 2314050005743 to N.J.), the Shenzhen Government ‘Peacock Plan’ (Y01416126 to N.J.), the Guangdong Provincial Science and Technology Innovation Council Grant (2017B030301018 to N.J.).

## Author contributions

J.T.Z. and Y.X. undertook biochemical studies, from sample preparation and purification, biochemical assays. J.T.Z also performed cryo-EM data collection data processing, and structure refinement. N.C. contributed to the cryo-EM sample preparation, and R. T. participated in the helpful discussions regarding this project. N.J. directed the research. N.J. and J.T.Z. wrote the manuscript with input from other authors.

---

**Extended Data Fig. 1.**
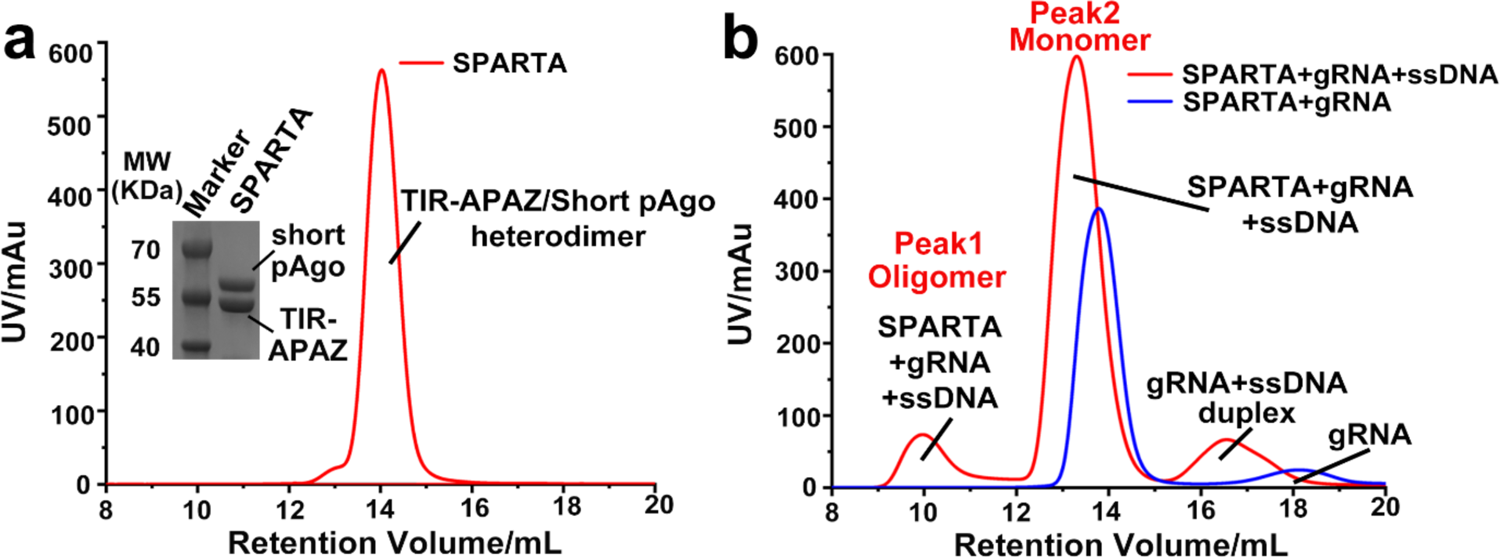
*In vitro* assembly of the SPARTA^gRNA^-ssDNA target complex. (a) Size exclusion chromatography and SDS-PAGE profiles of the purified SPARTA complex. (b) Size exclusion chromatography profile of the SPARTA^gRNA^ and SPARTA^gRNA^-ssDNA complexes.

**Extended Data Fig. 2.**
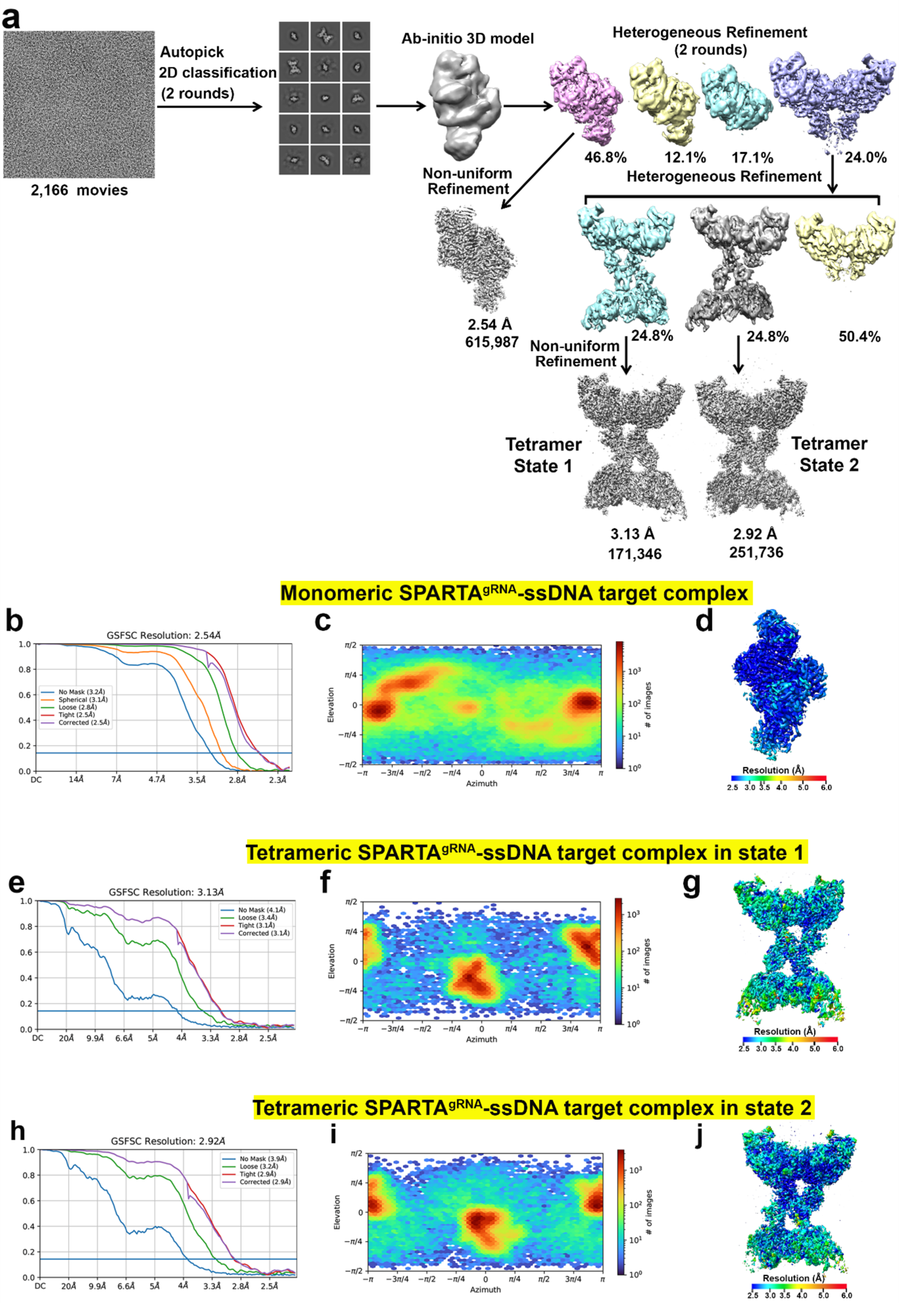
CryoEM reconstruction of monomeric and tetrameric SPARTA complexes in two different assembly states. (a) Flow chart of image processing for SPARTA complexes. (b, e and h) Fourier Shell Correlation curve of the monomeric SPARTA complex (b) and the tetrameric SPARTA complex in assembly state 1(e) and state 2 (h). (c, f and i) Direction distribution plot of the monomeric SPARTA complex (c) and the tetrameric SPARTA complex in assembly state 1 (f) and state 2 (i). (d, g and j) Final three-dimensional reconstructed map of the monomeric SPARTA complex (d) and the tetrameric SPARTA complex in assembly state 1 (g) and state 2 (j), colored according to local resolution.

**Extended Data Fig. 3.**
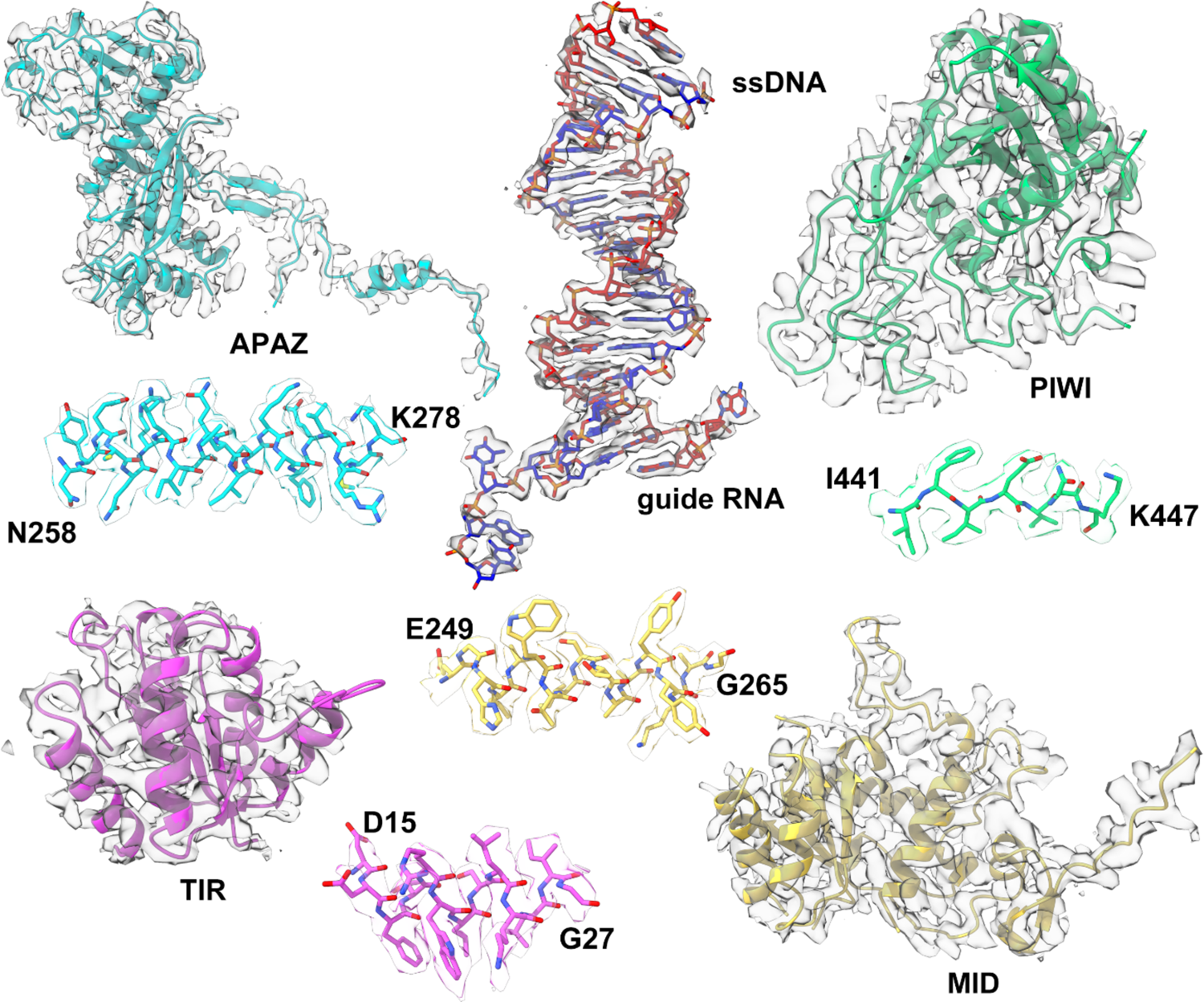
CryoEM densities of each domain and the gRNA-ssDNA target duplex in the monomeric SPARTA^gRNA^-target ssDNA complex structure. Densities for indicated regions are shown in the context of the atomic model.

**Extended Data Fig. 4.**
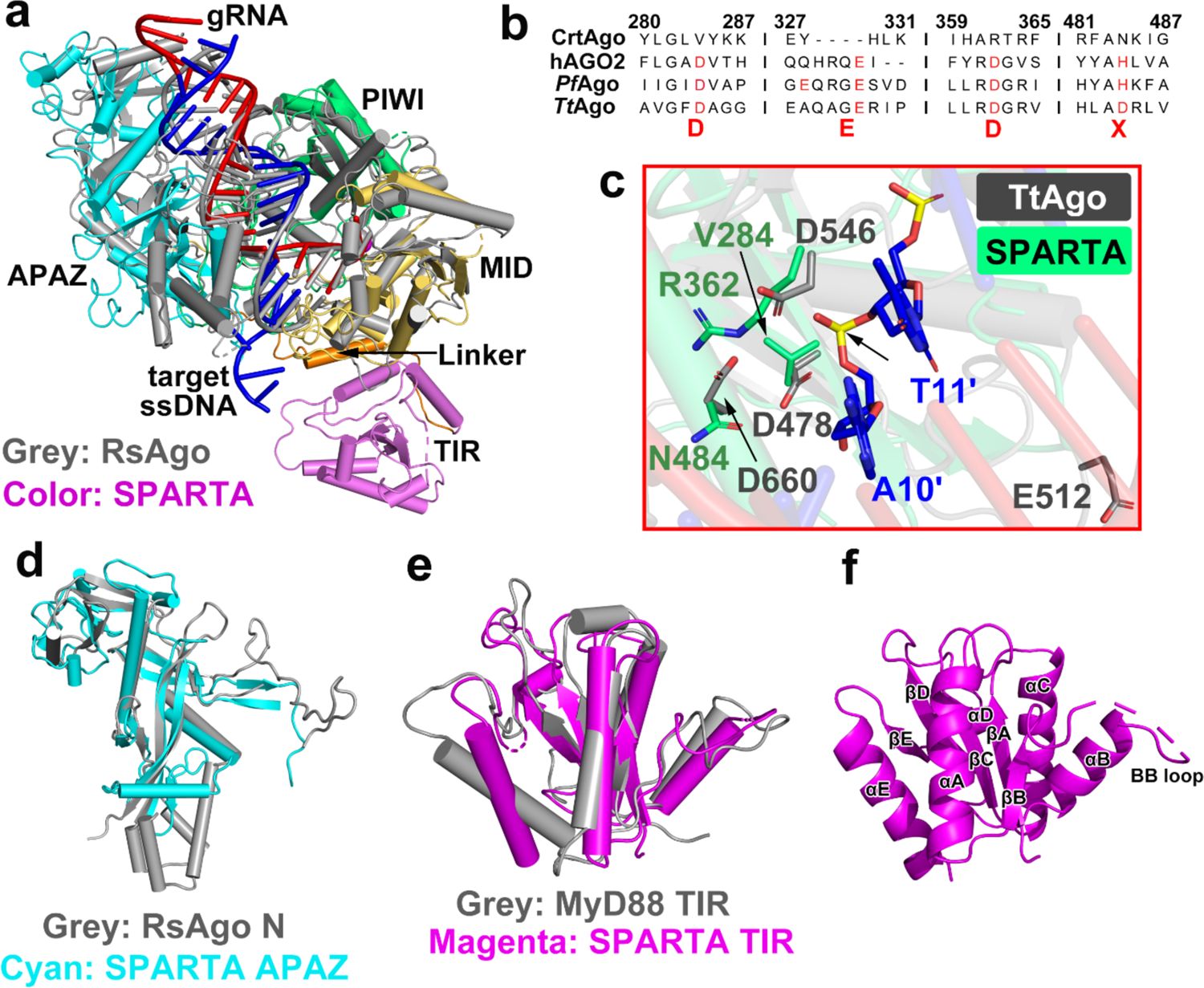
Structure and sequence comparisons of the SPARTA complex with related proteins. (a) Structural comparison of the prokaryotic *R. sphaeroides* RsAgo complex (PDB 5AWH, in grey) and monomeric SPARTA^gRNA^-ssDNA complex (in color). (b) Multiple sequence alignment of the conserved catalytic tetrad motif in the PIWI domain of *C. thermophila* SPARTA, NP_036286.2 (*Homo sapiens* AGO2), WP_011011654.1 (*Pyrococcus furiosus* Argonaute) and WP_011174533.1 (*Thermus thermophilus* Argonaute). The catalytic motif DEDX is indicated with red font. (c) Structural comparison of the catalytic tetrad between *T. thermophilus* TtAgo (grey, PDB 3F73) and *C. thermophila* SPARTA (dark green). (d) Structural alignment of the *R. sphaeroides* RsAgo N domain (grey, PDB 5AWH) and *C. thermophila* SPARTA APAZ domain (cyan). (e) The *C. thermophila* SPARTA TIR domain (magenta) structurally resembles the *H. sapiens* MyD88 TIR domain (grey, PDB 7BEQ). (f) Ribbon representation of the SPARTA TIR domain. The conserved structural elements (αA– αE, βA–βE) and the BB loop critical for NADase activity are labeled.

**Extended Data Fig. 5.**
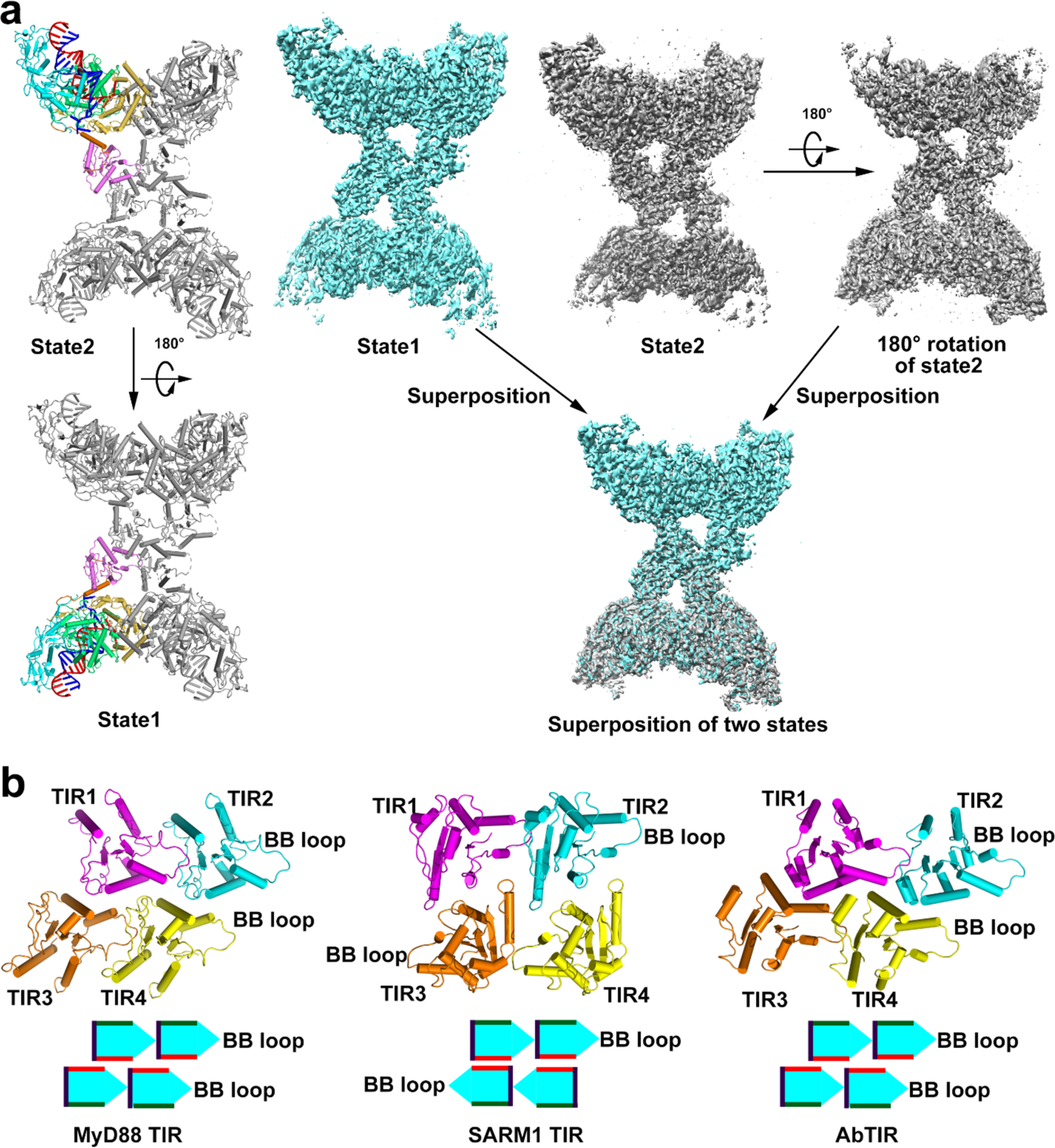
TIR domain assembly patterns. (a) Ribbon (left) and surface (right) comparision of tetrameric SPARTA complex in two different states. Tetrameric SPARTA in the second state can transfer to the first state via 180° rotation horizontally. (b) Ribbon and schematic representations of TIR domain assembly patterns in *H. sapiens* MyD88 (PDB 7BEQ), *H. sapiens* SARM1 (PDB 6O0Q) and AbTIR (PDB 7UXU).

**Extended Data Fig. 6.**
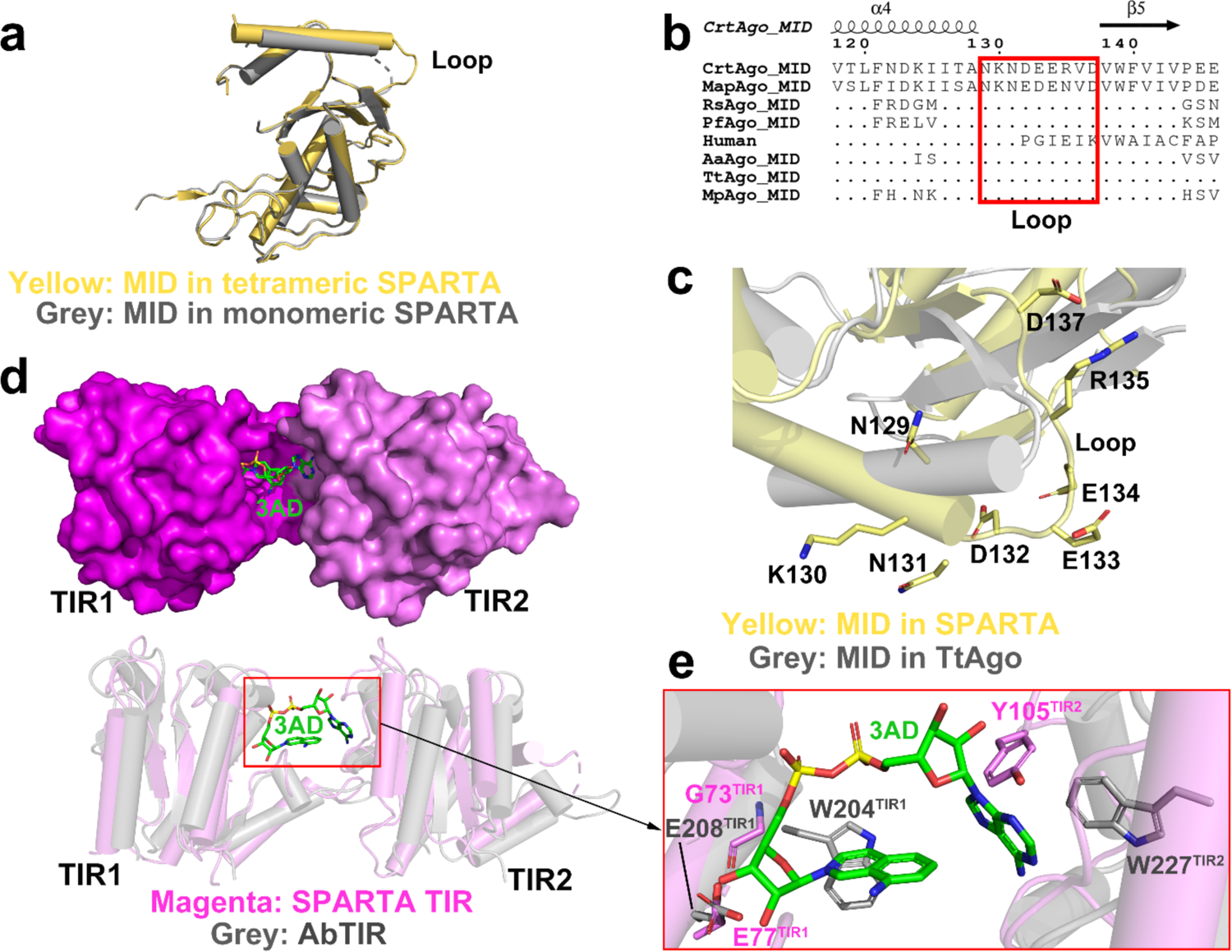
Structural comparison of the MID and TIR domains of SPARTA and other proteins. (a) Structural comparison of the MID domains of the monomeric SPARTA complex (grey) and the tetrameric SPARTA complex (yellow). (b) Multiple sequence alignment of the loop in the SPARTA MID domain and other Agos. The aligned sequences are from WP_109649955.1 (*Maribacter polysiphoniae* Argonaute, MapAgo), ABP72561.1 (*Rhodobacter sphaeroides* Argonaute, RsAgo), WP_011011654.1 (*Pyrococcus furiosus* Argonaute, PfAgo), NP_036286.2 (human AGO2), WP_010880937.1 (*Aquifex aeolicus* Argonaute, AaAgo), WP_011174533.1 (*Thermus thermophilus* Argonaute, TtAgo), WP_014295921.1 (*Marinitoga piezophila* Argonaute, MpAgo). (c) Structural alignment of the loop in the SPARTA MID domain with *T. thermophilus* Ago (grey, PDB 3HVR). (d) Docking of NAD^+^ into the TIR domains of tetrameric SPARTA, and superimposition of SPARTA TIRs with AbTIR (PDB 7UXU). (e) The putative NAD^+^ binding pocket in the SPARTA TIR domains superimposed with AbTIR.

**Extended Data Fig. 7.**
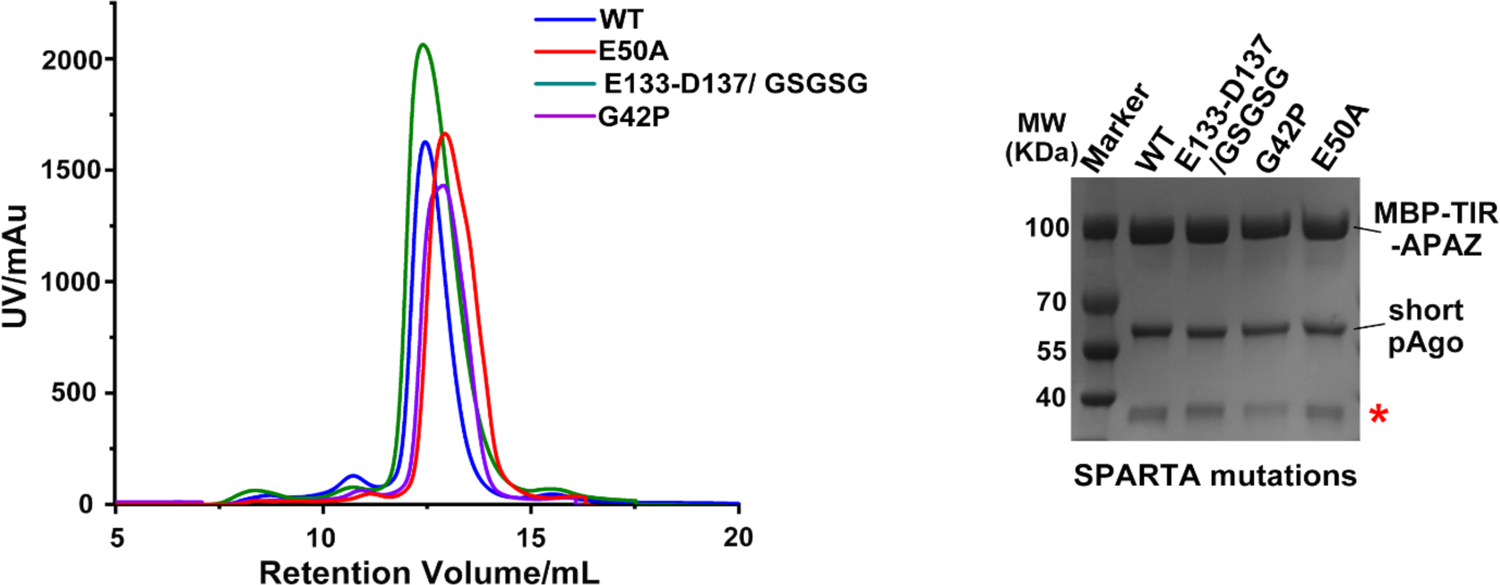
Size exclusion chromatography analysis of SPARTA mutants. Size exclusion chromatography (left) and SDS-PAGE profiles (right) of SPARTA mutants. E133-D137 (GSGSG) indicates the replacement of loop (133-EERVD-137) with a GSGSG linker in the MID domain. G42P and E50A indicate substitution of residues G42 and E50 in the TIR domain into proline and alanine, respectively. A MBP tag is fused to the N terminal of TIR-APAZ subunit in all SPARTA mutants. Red star indicates the individual MBP tag.

**Extended DataTable 1.**
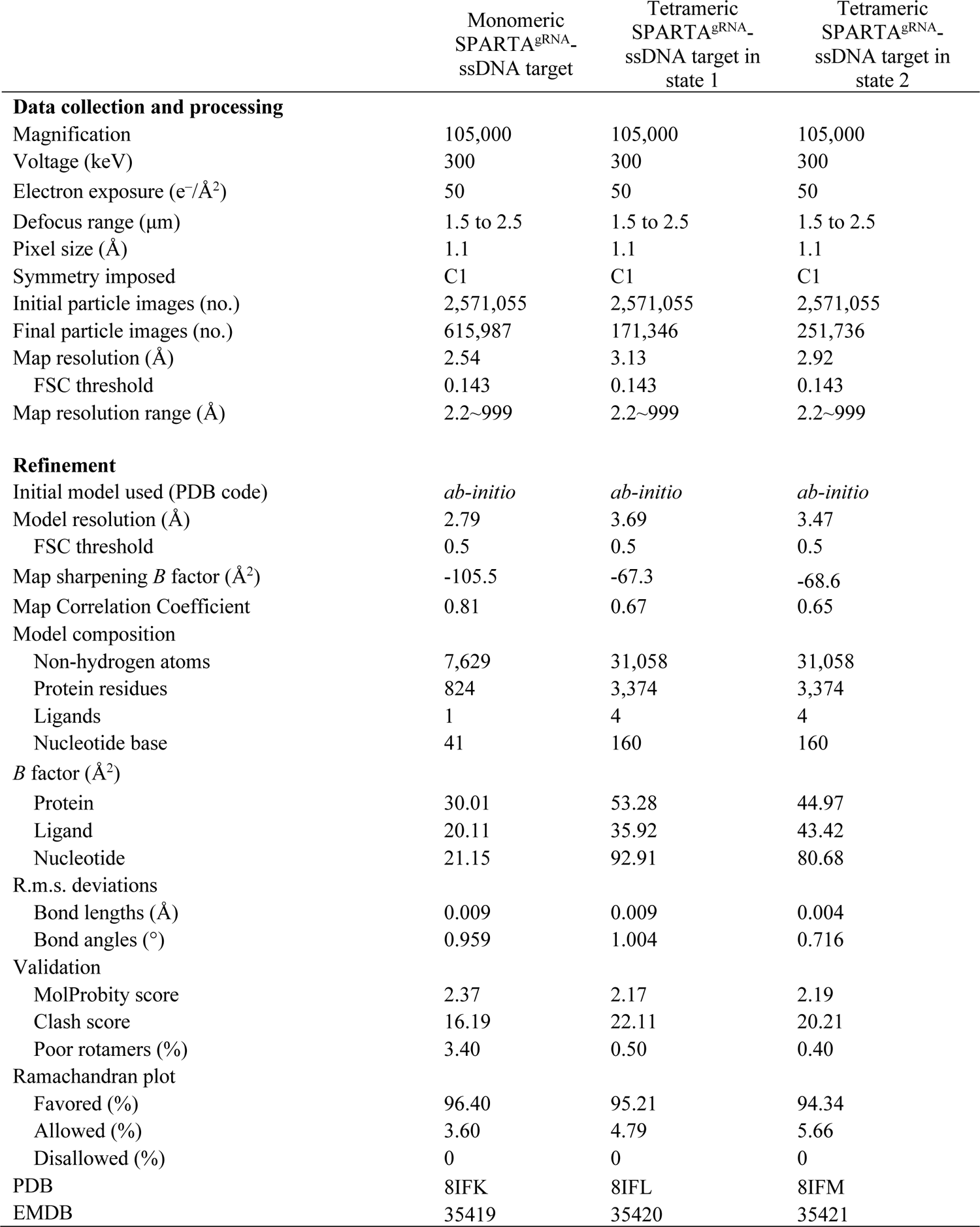
Cryo-EM data collection, refinement and validation statistics

